# Thermodynamics and free energy landscape of BAR-domain dimerization from molecular simulations

**DOI:** 10.1101/2020.12.09.418020

**Authors:** Adip Jhaveri, Dhruw Maisuria, Matthew Varga, Dariush Mohammadyani, Margaret E Johnson

## Abstract

Nearly all proteins interact specifically with other proteins, often forming reversible bound structures whose stability is critical to function. Proteins with BAR domains function to bind to, bend, and remodel biological membranes, where the dimerization of BAR domains is a key step in this function. Here we characterize the binding thermodynamics of homodimerization between the LSP1 BAR domain proteins in solution, using Molecular Dynamics (MD) simulations. By combining the MARTINI coarse-grained protein models with enhanced sampling through metadynamics, we construct a two-dimensional free energy surface quantifying the bound versus unbound ensembles as a function of two distance variables. Our simulations portray a heterogeneous and extraordinarily stable bound ensemble for these modeled LSP1 proteins. The proper crystal structure dimer has a large hydrophobic interface that is part of a stable minima on the free energy surface, which is enthalpically the minima of all bound structures. However, we also find several other stable nonspecific dimers with comparable free energies to the specific dimer. Through structure-based clustering of these bound structures, we find that some of these ‘nonspecific’ contacts involve extended tail regions that help stabilize the higher-order oligomers formed by BAR-domains, contacts that are separated from the homodimer interface. We find that the known membrane-binding residues of the LSP1 proteins rarely participate in any of the bound interfaces, but that both patches of residues are aligned to interact with the membrane in the specific dimer. Hence, we would expect a strong selection of the specific dimer in binding to the membrane. The effect of a 100mM NaCl buffer reduces the relative stability of nonspecific dimers compared to the specific dimer, indicating that it would help prevent aggregation of the proteins. With these results, we provide the first free energy characterization of interaction pathways in this important class of membrane sculpting domains, revealing a variety of interfacial contacts outside of the specific dimer that may help stabilize its oligomeric assemblies.

## I. Introduction

In the cell, reversible protein interactions allow for dynamic signal transduction and controlled nucleation and remodeling of multi-protein assemblies, such as occurs during the membrane remodeling processes of endocytosis^1^ and eisosome formation^2^. The Bin-Amphiphysin-Rvs (BAR) domain family are highly structured banana shaped domains that contain interfaces both for protein-membrane interactions, and protein-protein interactions with other BAR domains^3^. The formation of the hetero/homodimers of BAR containing proteins is a key step supporting their function as membrane binders and sculptors. To induce membrane bending and tubulation, the specific BAR dimers form additional protein-protein contacts, assembling into higher-order oligomers on the membrane surface^4^, and in some cases in solution^4^. The efficiency of formation and stability of this higher-order assembly will vary depending on the stability of individual BAR domain homo/heterodimers. While the solution dimers of various BAR domains and the membrane bound oligomers have been characterized structurally through experiment^2–9^ and molecular simulations^10–12^, the thermodynamics or kinetics of the BAR dimerization has received characterization only via experiment^5,13–15^. Here we use microseconds long, coarse-grained Molecular Dynamics (MD) simulations to quantify the free energy surface and binding affinity of BAR domain homodimerization in solution. Combining molecular simulations and enhanced sampling with metadynamics^16^, we can characterize the full bound and unbound equilibrium ensembles, providing a mechanistic understanding of the dimerization process and a foundation for future comparison with other BAR sequences or under changes to solution and membrane conditions.

BAR domain containing proteins have been a focus of broad structural, functional and biochemical characterization due to their central role in membrane remodeling, with critical roles in vesicle formation, actin dynamics and signaling^7,17^. The BAR domain superfamily of proteins consists of the classical BAR, N-BAR(N-terminal amphiphatic helix), F-BAR (Fer/CIP4 homology-BAR) and I-BAR (IRSp53 MIM homology-BAR). They all form rigid dimers that impart an intrinsic curvature to the membrane, but have variations in structure that induce or sense distinct shapes of curvature^18^. The dimerization of most BAR proteins leads to a concentration of positive charges on its concave surface that predominantly forms the membrane binding interface^19,20^. The weak sequence homology across different BAR domains indicates that each BAR dimer could be stabilized by different intermolecular interactions. For endophilin A1, mutation of a single hydrophobic residue, located at the BAR domain interface, can impair dimerization^21^, whereas for the SNX33 BAR domain interface, several amino acids provide specificity to the dimer contacts^22^. Experimental studies have reported dimerization affinities in solution in the range of sub-nanomolar to micromolar, leading to different mechanistic explanations^15,20,21^. The FCho2 BAR dimers bind at 2.5uM^5^, with a similarly moderate value of 10uM measured for endophilin^20^. In contrast, a very strong value of 0.5nM was also reported for endophilin^15^. This demonstrates the sensitivity of the dimer stability to the sequence and structure of the monomers, which will ultimately determine their function *in vivo*.

The yeast BAR domain protein LSP1, which we study here (Fig 1), plays an essential role in membrane remodeling via eisosome formation, along with a homologous BAR domain PIL1^4 2^. LSP1 and PIL1 are structurally related to the classical BAR proteins endophilin 2, amphiphysin, and arfaptin2, but have minimal sequence homology to them^2^. The LSP1 homodimer is stabilized by an enrichment of hydrophobic interactions between the inner surface of the monomers, with a buried surface area of ~6000 A^2^ that is relatively large for protein-protein association^23^. LSP1 forms dimers that further oligomerize into stable helical filaments *in vitro* in solution, without even requiring localization to the membrane, indicating relatively stable protein-protein contacts that can form simultaneously with the crystal dimer^4^. LSP1 also forms oligomeric scaffolds on the plasma membrane of yeast, along with PIL1^2^. Despite 72% sequence homology, PIL1 differs functionally from LSP1, forming more diverse and stable multimeric complexes *in vitro* ^4^ *⍰*, and resisting crystallization, unlike LSP1^2,4^.

**Figure 1.**
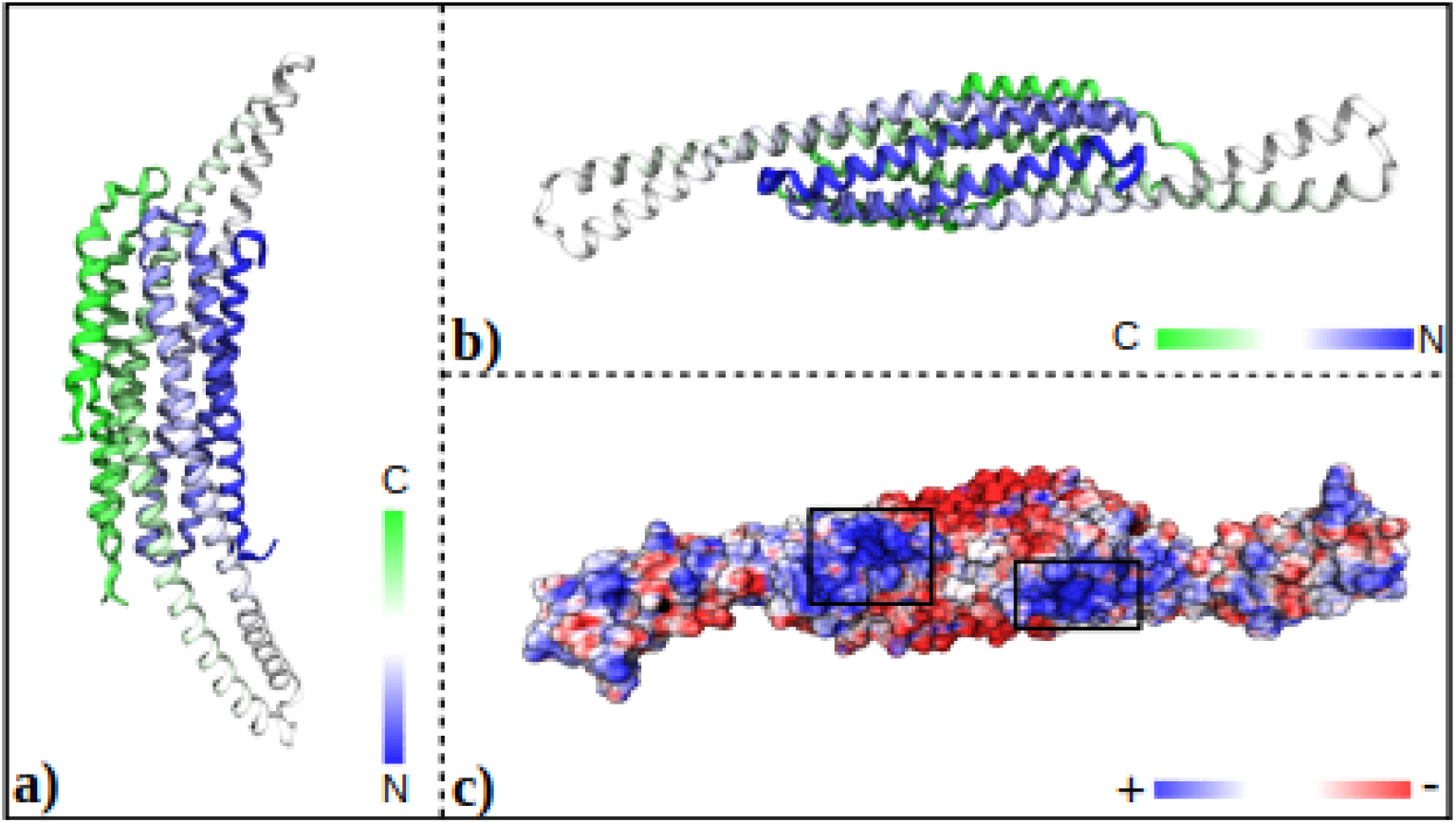
a) Crystal structure of the LSP1 homodimer, showing both chains A and B represented in a ribbon form. Each chain follows a gradient from blue (N terminus) to green (C terminus). b) LSP1 homodimer rotated by 90° around the z-axis and 90° again around the x-axis relative to a. The color scheme has is same as in a. c) Surface representation of the homodimer colored according to electrostatic potential and in similar orientation as in b. We can see positively charged, blue colored, patches (highlighted by black boxes) of the dimer, that interacts with the negatively charged membrane.

MD simulations offer an attractive alternative to study the dimerization of BAR domain proteins, because it obtains high-resolution structural and thermodynamic information simultaneously. However, such studies are computationally demanding. Measurements of the binding thermodynamics of any protein-protein binding pairs from simulation are still rare. Unlike protein interactions with small molecules, protein-protein association involves relatively large surface areas that can form a variety of nonspecific encounter complexes that may contribute to formation of the bound structure^24^⍰. Binding free energies can be estimated from well-defined bound and unbound configurations sampled from MD^25,26^⍰, but reconstructing a free energy surface to predict the ensemble of bound and encounter complexes along reaction coordinates, as we do here, requires multiple transitions between the bound and unbound states. Using coarse-grained models of proteins, Monte Carlo simulations^24,27^ ⍰ and Brownian dynamics simulations^28^ have been used to successfully characterize bound ensembles or equilibrium dissociation constants. Atomistic MD simulations of protein-protein association have been recently applied to measure binding kinetics^29,30^ and binding free energies of the barnase-barstar interaction^29,31^, and binding pathways of four additional structured pairs⍰31. Here we perform MD simulations with the coarse-grained MARTINI force field^32^, which was recently used to study binding free energies of transmembrane helix dimerization^33^. MARTINI provides a significant speed up in computer processing time without dramatic reduction in molecular resolution, retaining explicit solvent and flexible (not rigid) body dynamics. For association, a relatively large box size is necessary to prevent monomers from interacting through multiple images, which unavoidably increases the simulation cost. For all protein-protein association studies using MD (all-atom or the coarse-grained model we use here), thermodynamics and binding pathways are extracted in combination with enhanced sampling techniques.

Several enhanced sampling techniques have been used to extract binding free energies and free energy surfaces from MD simulations, including umbrella sampling^34^⍰, tempered binding⍰31⍰, replica exchange^24,27,35^⍰, Markov state models^29^, simulated annealing^36^, weighted ensemble (WE) protocols^30,37^ and metadynamics^33,38,39^. Free energy differences are calculated from Boltzmann weighted ensemble averages of atomic configurations corresponding to each state. Free energy perturbation^40^and thermodynamic integration^41^are primarily applied to quantifying interactions between proteins and ligands that have a single bound state, rather than large binding surfaces sampled in protein-protein association. Free energy calculations are sensitive to (i) Choice of a suitable force-field (ii) treatment of long-range interactions (iii) sampling of all relevant states and (iv) convergence^42,43^. MARTINI offers a well-studied force-field (i) for both proteins and membranes, and includes long-range interactions (ii) and explicit solvent. For enhanced sampling, which ⍰helps achieve (iii) and (iv), we use metadynamics^16^. A primary reason behind using metadynamics is to characterize the range of possible protein-protein complexes formed, allowing us to predict which is most stable, and quantify the relative stability of nonspecific dimers. Particularly for oligomer forming proteins like LSP1, this allows us to assess whether interfaces outside of the specific dimer are sterically accessible to form between two pairs of dimers. Further, because a protein-protein interface is much larger than a protein-ligand binding pocket, at least two reaction coordinates or collective variables (CVs) are needed to distinguish distinct bound configurations. Metadynamics is efficient even for constructing multidimensional free energy surfaces (FES)^44,45^⍰. The FES identifies intermediate states, barriers, and transition pathways^46,47^⍰, rather than just calculating a free energy difference between two reference states, and metadynamics includes well-controlled strategies to improve convergence^48–50^ and quantify unbiased thermodynamic quantities.

In this study, we first describe how we can extract free energies, entropies, and enthalpies from our simulations, and classify bound sub-ensembles using structure-based clustering. We quantify the dependence of our standard Gibb’s free energies of binding (or dissociation constants K_D_) on the definitions of bound vs unbound ensembles, due to the significant contributions of nonspecific complexes to the state space. We evaluate the convergence and robustness of our free energy surfaces with changes to the metadynamics parameters. We show that the addition of 100mM NaCl causes small but important shifts in our bound state ensembles, suggesting a significant role of the solvent. We discuss the over-stabilization of our bound ensemble, which at ~90k_B_T is more similar to protein folding energies than protein association. Lastly, we discuss the implications of the heterogeneous bound state ensemble we find for these modeled LSP1 homodimers. Our diverse bound ensemble supports contacts outside of the specific dimer that could stabilize the oligomer. However, the relative depths of these minima on the FES also indicates some limitations of the force-field used, as the crystal structure conformation is not significantly more probable or stable than nonspecific associations without salt added, which would render aggregation likely and crystallization difficult. Overall, however, our results provide a quantitative characterization of BAR domain dimerization that assesses structural, thermodynamic, and solvent contributions to the variety of protein-protein complexes formed by the pair of monomers.

## II. METHODS

### II.A Calculations of thermodynamic quantities

In a protein dimerization reaction: *L + L* ⇌ *LL* the equilibrium dissociation constant is related to the standard Gibbs free energy of binding by the following expression^51,52^

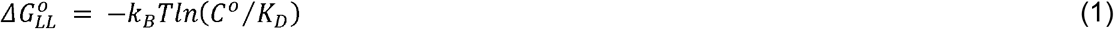

Where k_B_ is the Boltzmann constant, T is the temperature, *C°* is the standard state concentration of 1M, and ΔG°_LL_ is the standard Gibb’s free energy difference between the bound (LL) state and the unbound (L) state for a system at constant N, P, and T. To evaluate the 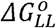, we use the following expression^51^

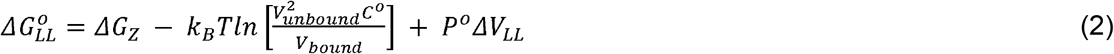

Where V_unbound_, V_bound_ are the simulation volumes of the unbound and bound states respectively, P° is the pressure, and *ΔV*_*LL*_ is the change in volume between the bound and unbound ensembles. Here, the second term represents the standard state correction ⍰. The standard state correction for our simulation volume is 20kJ/mol, or 8k_B_T. The volume correction (third term) is negligible. We calculate *ΔG*_*z*_ from the canonical partition functions of the bound and the unbound ensembles, which can be expressed as -

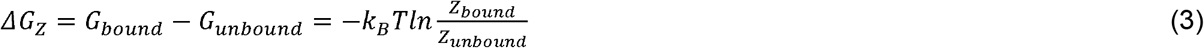

The partition functions are calculated by integrating over the Boltzmann weighted free energies of the bound or unbound ensembles such that

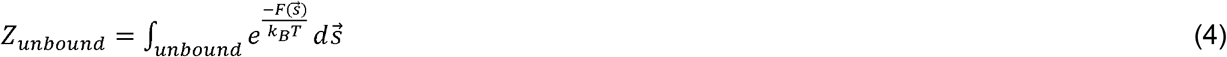

and similar for the bound state, where 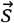 are the variables 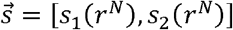 used as reaction coordinate to collect the free energy surface 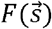 and r^N^ are the microscopic coordinates of the N atoms/beads. The values of *ΔG*_*z*_ thus depends on the definition of bound and unbound ensembles we integrate over, which we quantify in the results. We construct the free energy surface 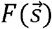 from performing MD simulations with metadynamics biasing, as described below.

The entropy change between the bound and unbound ensembles can then be evaluated by^52^

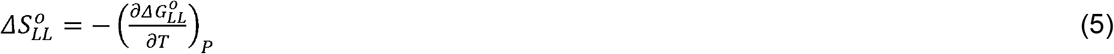

When applied to Eq 2, this gives

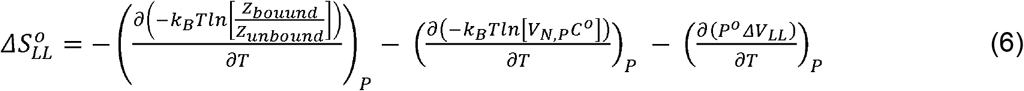

which leads to:

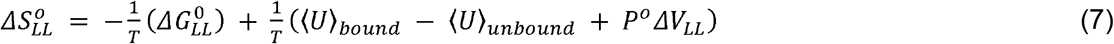

where *U* is the potential energy, and we assume the average kinetic energy is the same between bound and unbound states. We can define the average potential energy for any sub-ensemble Ω using -

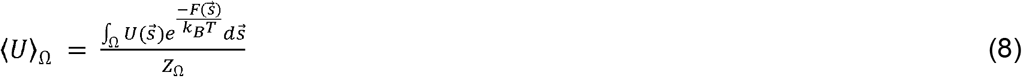

with the denominator defined in Eq 4 and 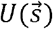 is the potential energy averaged over all the microscopic configurations belonging to that region in CV space, (in our case a particular grid point) and is given by the expression:

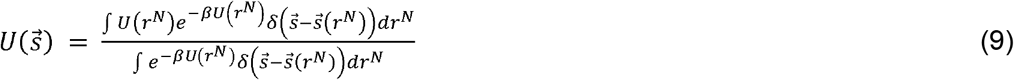

We discuss below the correction used when sampling is biased.

### II.B Calculating the free energy surface from simulations with metadynamics

#### Metadynamics background

We briefly summarize here the metadynamics approach, which has been extensively described elsewhere^44^. Metadynamics biases the system along a reduced set of degrees of freedom called collective variables (CVs), 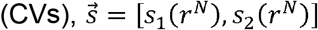, where here we use two. The CVs are a function of the atomic coordinates and are chosen by the user to best distinguish the coarse states of interest, in our case the bound and unbound states. Throughout the MD simulations, a history dependent potential term (called a bias potential), is accumulated along the CVs and added to the Hamiltonian of the system to drive the dynamics.

The bias potential is given by

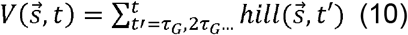

where 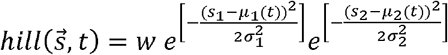, the hill means *μ*_*i*_(*t*) are given by the instantaneous values of the *i*^th^ CV at time *t*, *τ*_*G*_ is the time interval for adding each hill (Gaussian bias potential), *w* is the weight of the unnormalized Gaussian and *σ*s are the width of them for each CV. After sufficient time *t*, the bias potential will converge to the unbiased free energy surface modulo a constant offset^16^:

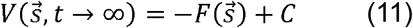

which is the FES used in the thermodynamic calculations described above, as

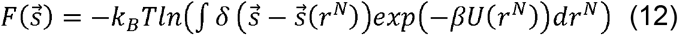

The PLUMED plugin^53^ performs the biasing along our simulation trajectory.

#### Choice of coarse variables

We constructed two-dimensional free energy surfaces along a pair of CVs. To differentiate the bound and the unbound states, we used two distance CVs as shown in Figure 2a. The distance d_1_ is calculated between the center of geometry (COG) between the patch on monomer A containing residues 57LYS – 64THR, and the COG of the patch on monomer B containing residues 78GLU-92ASP. The distance d_2_ is calculated between the COG of the patch on monomer A containing 224LEU-229ASP and the COG of the patch on monomer B containing residues 189ASN-196LYS. These patches were chosen because they are in contact in the bound state, defined via the crystal structure 3PLT. They are also at either end of the bound dimer, which allows us to distinguish distinct orientations, which would not be possible using just a single distance variable.

**Figure 2.**
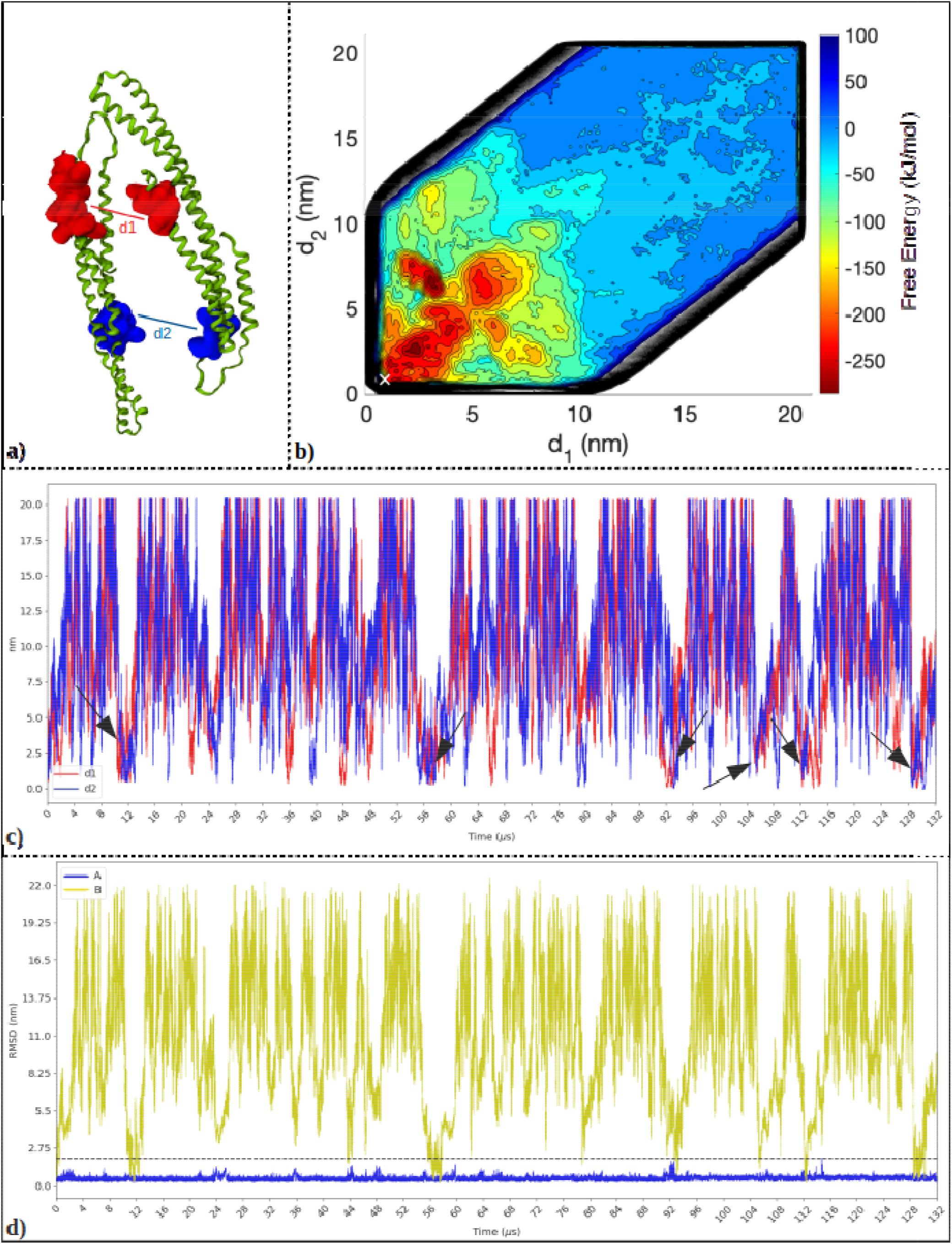
a) LSP1 chains A and B with the CVs, d_1_ and d_2_ between the two binding sites. (shown as surface representation in red and blue). b) Contour plot of the free energy as a function of d_1_ and d_2_ as calculated from metadynamics. This is the instantaneous FES after 132 μs. The ‘**X**’ represents the CV values for the crystal structure. c) Progression of d_1_ (red) and d_2_ (blue) with time. The arrows represent transition events to the native bound state, calculated where the RMSD between our dimer and the crystal structure drops below 2 nm.

#### Choice of metadynamics parameters

To perform metadynamics biasing of our simulations^53^, we had to select values for the Gaussian widths σ, the deposition time *τ*_*G*_, and the weights *w*. The widths were chosen along with the deposition times to provide sufficient resolution of the free energy surface and efficient sampling. We thus used a width of 0.1nm for both distances d_1_ and d_2_. We calculated the deposition time *τ*_*G*_ for applying a new gaussian bias along both coordinates based on an average relaxation time of d_1_ and d_2_. We performed short unbiased simulations to evaluate the average displacements of d_1_ and d_2_, when in the bound state. The time interval that gives rise to a distribution of displacements with width σ=0.1nm was used to define *τ*_*G*_. We used a value of *τ*_*G*_ = 140*ps*, and as a more conservative verification, we also used values of *τ*_*G*_=500ps. We did not see significant differences when using the faster time deposition, which showed faster convergence. For the weights or heights of the unnormalized gaussians, we ran the first ~8us of the simulations with a value of 10kJ/mol, which provided a balance of efficient biasing with sufficient resolution of the surface topology. We then decreased the height to 5kJ/mol for the remainder of the simulations. We did not use any well-tempered form of metadynamics, as we did not know in advance what the size of the free energy barriers would be. For the 100mM NaCl simulations, we used *τ*_*G*_ = 500ps, a height of 10kJ/mol, and widths of σ=0.1, 0.1nm.

### II.C Numerical evaluation of energies

#### Partition functions

To evaluate Eq 3 from our simulations, we need to integrate over the Boltzmann weighted free energies. We do this by representing the surface on a grid, assigning each grid point to a specific ensemble, and then numerically integrating over these grids. Grid points to define each sub-ensemble were defined by making cutoffs in the CV space depending upon the free energy surface. The free energy or potential energy of any sub-ensemble was reported as a *Δ*G by comparison to the unbound ensemble we defined as all grid points where d_1_ >12.5 and d_2_ >12.5. These distances were chosen based on where the contours of the FES flattened out. We also tested the free energy values found when we classified each configuration in our trajectory based on its microscopic properties. For example, we performed one calculation where the bound state was defined as the ensemble of configurations where the two chains are in contact. A contact was defined when the minimum distance between any residue pair of the two monomers was less than 0.8 nm. The corresponding grid points for each state were then defined by binning each configuration based on their d_1_,d_2_ values. However, we found these distance-based state definitions were not reliable determinants of the bound and unbound ensemble due to the heterogeneity of configurations at each grid point.

#### One dimensional (1D) free energy

To evaluate a 1D FES from our 2D FES, we re-calculate the partition function integrating over all values of d_1_ or d_2_. Specifically, 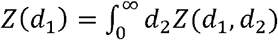. Then *G*(*d*_1_) = −*k*_*B*_*Tln(Z(d*_1_)).

#### Potential energies from biased simulations

To calculate 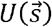 from an equilibrium NPT trajectory, one would normalize exploit the ergodic principle to take a simple unweighted time average over the potential energy of configurations within each grid point, 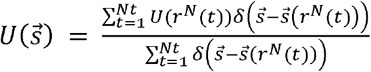 the total number of configurations from a trajectory. However, because we are performing biased sampling via the potential *V*(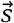, t) added to our Hamiltonian, each configuration does not appear according to its Boltzmann weight, and thus we must apply a reweighting factor as we average, giving

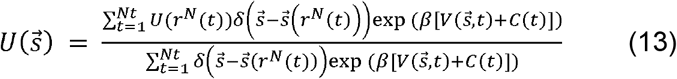

using the algorithm given in Tiwary and Parinello^54^, where *C*(*t*) is a constant offset to the potential function, which increases with increasing simulation time.

#### Average FES calculation

Once the basins on the free energy surface have been filled via the metadynamics sampling, the bias potential will continue to accumulate as the collective variables diffuse through the full space of CVs. Because there is a timescale of diffusion in CV space, this results in asymmetric sampling of one region of CV space, even after the free energy surface has been ‘flattened’, at a time *T*_*D*_. As a result, one observes oscillations in the FES on the timescale of the diffusion throughout the basins on the FES. As we see below, the diffusion along the distance CVs is relatively slow in the unbound state, resulting in slow oscillations of ~20-40*μ*s. We use one approach of averaging the FES after a time TD, which if the simulation is now truly performing a diffusive search in CV space, should provide a converging estimate of the time-independent FES. Based on the number of transitions in and out of the bound state (4) by 72*μ*s, and also the clear oscillations in measurements of ΔG, we use the surface at *T*_*D*_=72*μ*s as a starting point for calculating an average FES. Using the equation^16^

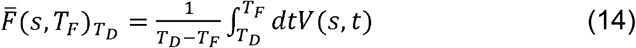

where *T*_F_ is the final time point, one can show that this is equivalent to applying a dampening weight to each gaussian potential that is deposited after T_D_ by rewriting the above using eq. 10 to get:

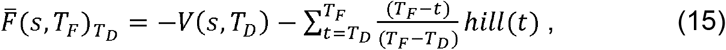

Where *hill(t)* is the gaussian deposited at time t. This thus effectively dampens the oscillations that occur as the sampling continues, giving the hills up to *T*_*D*_ a unitary weight to the FES based on the assumption that they are filling up meaningful basins of the surface, with later ones after *T*_*D*_ as time progresses more likely to contribute to oscillations on the FES height.

#### Error estimates

The error was estimated by putting the time-dependent values of an observable, such as ΔG(t), into 2-3 chunks, where the number of chunks *M* was decided based on removing correlation between adjacent chunks. We took the standard deviation σ of the means calculated for each chunk, to report the 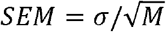.

### II.D Simulation Details

#### System setup

The LSP1 structure was obtained from the PDB under accession code 3PLT^2^ (Fig 1). To study the dimerization process, chains A and B were used. Chain B has a stretch of missing residues, that we modeled back in using the structure of chain A. The shortened form of the full LSP1 sequence was crystallized, and only those residues are within our model. They were shown to still bind to the plasma membrane despite lacking the initial 51 and final residues^2^. The simulations were performed using the MARTINI force-field version 2.2 ^55^. For MARTINI, the protein structure was coarse-grained using the ELNEDIN model, which is a combination of an Elastic Network model (EN) with the MARTINI force field⍰^56^. This model integrates a structure-based molecular description with a physics-based force field to study intermolecular interactions while maintaining the structural integrity of individual monomers. The CG structure was generated using the *martinize.py* script using the elnedyn forcefield option with the spring force constant as 500 kJ mol^−1^ nm^−2^ and cut-off as 0.9 nm.

#### Simulation parameters

The MD simulations under the MARTINI force field were performed using the GROMACS package^57^. The protein complex was solvated using the standard MARTINI water model. With a charge per LSP1 monomer of −12, for the pure water simulations the pair were neutralized using 24 Sodium ions (NA+) in a simulation box of dimensions ~24 nm × 24 nm × 24 nm. The solvated protein with the counter ions was minimized using the steepest descent algorithm. The system was then equilibrated for a total of 10 ns. First, the system was equilibrated under constant NVT conditions for 1ns using a timestep of 10 fs and then for 4 ns with a 20 fs timestep. The temperature was kept constant at 310 K using the velocity rescaling method with a time constant of 1 ps. This was followed by simulations under constant NPT for 1ns using a timestep of 10 fs and then for 4 ns with a 20 fs timestep. The pressure was kept constant at 1 bar using Berendsen coupling with a time constant of 12 ps. For the production runs, performed under NPT, the pressure coupling was shifted to Parrinello-Rahman while maintaining the integration time step of 20 fs. The bonds were constrained by the LINCS algorithm for all the simulations.

The simulations were performed under periodic boundary conditions and using the Verlet scheme to build the neighbor lists. The non-bonded interactions were cut-off at 1.2 nm. The coulombic interactions were modeled using the Reaction field with *∊*_*rf*_ =∞ and *∊*_*r*_ = 15 to account for electrostatic screening. The Lennard-Jones (LJ) potential was smoothly shifted to zero within the cut-off distance. With the PLUMED plugin for metadynamics^53^, we used the GRID option. For post-trajectory analysis, molecular coordinates were recorded every 200ps for each simulation. Analysis was performed in python using the MDAnalysis^58^⍰ package. For visualization, the coarse grained structures were converted to an all-atom representation using the pulchra algorithm^59^. The structures were visualized using the VMD tool^60^⍰.

#### 100mM NaCl system

For simulations with a bulk concentration of 100mM NaCl, we used 2710 beads of Na+ and 2686 of Cl−. We also increased the size of the box to 35.5 × 35.5 × 35.5nm. The smaller box is large enough to prevent any extended linear chain of BAR proteins to form. Nonetheless, we used a larger box because for a small subset of configurations, the two distance CVs would be biased against configurations where a protein was interacting with effectively two other proteins at once, on one side via its partner, and on the other side via its partner’s periodic image.

### II.E Numerical Analysis of MD trajectories

The RMSD of each dimer relative to the crystal structure was calculated by superimposing chain A over its crystal structure counterpart, and calculating the RMSD for the residues in the binding patches of chain B. To characterize our bound ensemble, we performed kmeans clustering of the trajectory based on the d_1_ and d_2_ values of each configuration, into 6 clusters. Only those configurations were considered were chain A and B are in contact with each other, and where d_1_+d_2_<10.2. So each grid point in CV space has a unique cluster assigned to it. Each cluster has about the same number of configurations in it.

#### Analysis of contacts and orientations formed between monomer pairs

The residues on each protein chain were grouped together into 11 patches for analyzing the different protein-protein interfaces formed in each cluster. Patches d_1_ and d_2_ correspond to the distance CV patches defined before. The d_c_ patch consists of residues 206-213 on chain A, and residues 206-213 on patch B. Residues are always numbered according to their 3PLT.pdb values. The native patch-1 (NP-1) consists of residues 187-205 of chain A, and residues 214-242 of chain B. NP-2 consists of residues 81-103 of chain A and 52-54, 57-62, 64, 65, 67-69, 72-77 of chain B. The non-native patch-1 (NNP-1) consists of residues 65, 67-69, and 72-80 of chain A and residues 93-105 of chain B. The NNP-2 comprises the set of residues 106-125, 127-129, 131, 132, and 134-150 from each chain. The NNP-3 consists of residues 214-224 of chain A and residues 197-205 on chain B. The NNP-4 consists of residues 230-264 of chain A and 243-264 of chain B. The tail region includes residues 151-186 of chain A and residues 151-188 of chain B. The M patch is the membrane binding surface on each chain^2^ which includes residues R56, K63, K66, R70, R126, K130, R133. The contact patches termed d_1_, d_2_, d_c_, NP-1, NP-2, are considered as part of the native interface since they include most of the residues which are in contact in the native crystal structure.

The orientation of the membrane binding patches with each other was evaluated using distance and angle measurements. The distance d_M_, is defined as the distance between the center of geometries (COG) of the membrane binding patches M on each chain. The orientation angle between the membrane binding patches on chains A and B is defined as 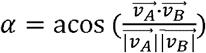, where 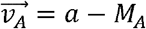, where *a* is the COG of the backbone beads of residues 193, 194, 198, 201 and M_A_ is the COG of patch M. 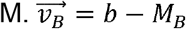, where *b* is the COG of backbone beads of residues 193, 217, 219, 224, 194, 200 and M_B_ is the COG of patch M. Another distance measurement, d_C_, is used to analyze the distribution of separations between the two chains. It is measured as the distance between the COGs of the d_C_ patches on each chain, as defined above.

## III Results

### III.A Free energy surface along two distance collective variables from metadynamics

We constructed the free energy surface (FES) using metadynamics along two distance collective variables (CV)s d_1_ and d_2_, shown in Fig 2a. The residues that define these patches are both part of the binding interface (Methods), and thus the distances are both very short in the crystal or bound state, with d_1_=0.818 nm, d_2_=0.883 nm. We chose patches at opposite ends of the binding interface to differentiate different dimer orientations. The accessible region of the FES is bounded by the physical inability to extend to a large separation in one CV if the other is in close contact, hence the diagonal shape (Fig 2b). After 132 μs of simulation, starting from the bound structure, we have observed multiple (6) transitions back into the natively bound state, where we denote the crystal structure by an ‘X’ on the FES (Fig 2b). The time evolution of d_1_ and d_2_ is plotted in Fig 2c. We define transitions to the natively bound state (black arrows) as occurring when the RMSD between our configuration and the crystal structure is <2nm. As expected, our FES shows an extended unbound region when both distance CVs extend beyond ~12nm. However, we also find several stable regions in CV space beyond the neighborhood of the crystal structure at the ‘X’. The bound state ensemble that we sample is thus quite large and diverse, and configurations with nonspecific interactions between the BAR dimers are comparable to the crystal structure in stability, particularly at d_1_=d_2_~4nm, and another region at d_1_~3nm and d_2_~6nm. As we characterize below, these structures form multiple residue contacts between the dimer, with both native crystal contacts and non-native residues participating. By integrating along either d_1_ or d_2_ in the calculation of the partition function (Methods), we can plot the 1D free energy landscape along either CV. We find that around values of ~12.5 in d_1_ and d_2_, the free energy flattens out relative to the bound structures. When considering the full bound state ensemble, we see dozens of transitions in and out of this region, but the minima adjacent to the crystal structure is not always sampled (see Movie S1).

### III.B Binding free energies from the Free energy surface

To obtain an accurate estimate of the binding free energy, the system has to exhaustively explore the CV space. However, due to the time required to sample large regions of CV space, accumulation of biasing potentials in, for example, the bound vs unbound ensembles produces a FES that temporarily overestimates the stability of that state. To evaluate convergence of our surface and our ΔG measurements, we quantify the impact of the time-dependent changes in the free energy surface upon free energy differences of specific sub-states, shown in Fig 4 (inset). We evaluate the partition function of each state by assigning regions or grid points in the CV space to bound and unbound states and then calculate ΔG using Eq 3. The unbound state is defined as the region outside the blue lines in Fig 4 (inset). For the bound state, we consider three regions, a specific bound state (defined by the black box), nonspecific bound state (defined by the magenta box) and a third, larger bound state, that lies inside the blue lines. The time-dependent free energy surface results in a time-dependent ΔG, shown in Fig 4a for each region. We see a slow oscillation of ΔG in time, which is what we expect if our surface is converging and is now re-visiting and refilling regions on the surface. We do observe that the magnitude of the oscillations is relatively large. Our bias potentials have a height of 5 kJ/mol after 8μs, so we expect oscillations of at least this size, but the timescales for traveling along our surface are also relatively slow, resulting in significant filling before a new region of space is visited. From ~84-100μs, for example, the system is exploring mostly unbound states, before it returns to visit different regions of the bound state. We can estimate an SEM by measuring ΔG over two uncorrelated ~30μs windows after 72μs (Methods). For the two-state model, we find ΔG =−242 ±22kJ/mol. For the specific bound state, we have ΔG =−196±39 kJ/mol, and the nonspecific state has ΔG= −214±10 kJ/mol.

The stability of the bound state can be attributed to not only the native state configurations but also multiple additional basins involving nonspecific contacts. Experimentally, a K_D_ is assumed to describe a two-state model of bound vs unbound states. Hence all configurations should contribute to either ensemble. For strong binders, there is often a clear distinction between the bound vs unbound states, as a shift in the cutoff between them will not change the ΔG^34^ ⍰ due to the Boltzmann weighting and averaging of the large values. However, we see from Fig 3 that there are configurations stretching out to >10nm that are clearly more stable than the unbound structures. We quantify how moving the boundary between the bound and unbound ensemble impacts the ΔG value (Fig 4b). The ΔG at our final time-point is stable after the 12.5nm cutoff, which is what we use for the unbound ensemble, although we note due to fluctuations of the ensemble, the mean value is still decreasing slightly beyond this cutoff (Fig 4b). We also calculated the ΔG after averaging the FES starting after 60μs or 72 μs, which amounts to adding in each subsequent bias potential after this time with an increasingly small weight (Methods). For the averaged surface, we find smaller ΔG values of −192kJ/mol and −174kJ/mol after 60μs or 72 μs. This is because none of the basins in the bound state are persistently the lowest energy regions on the surface, and thus averaging over time results in all bound state basins becoming slightly shallower.

**Figure 3.**
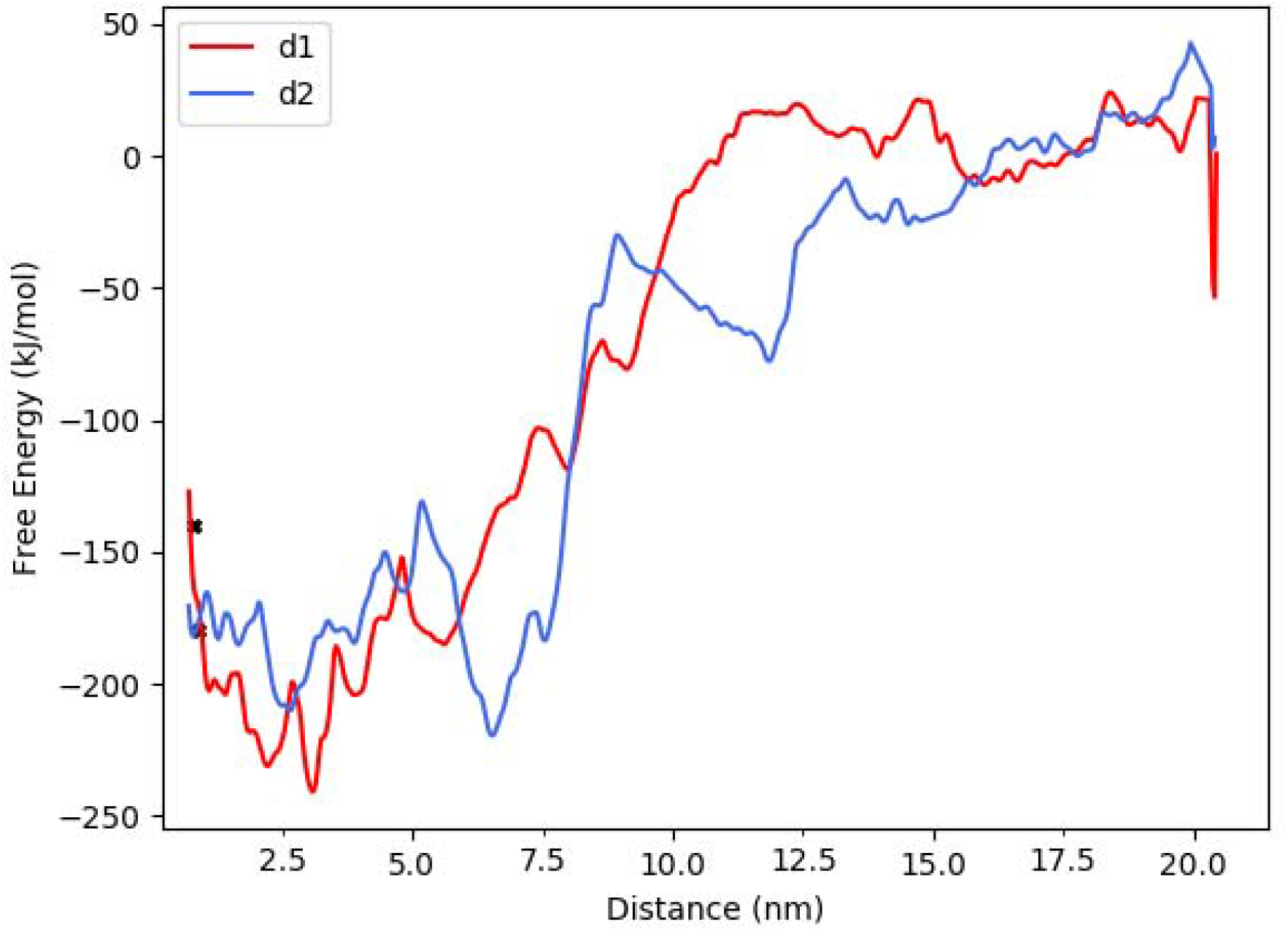
FES integrated along one CV only. In red, F(d_1_) and in blue F(d_2_). We re-zeroed the FES based on the values of free energy at d_1_ = d_2_ =17.5. The crystal structure location is shown in black x-marks for d_1_=0.818nm and d_2_=0.883nm. This shows the roughness of the free energy surface, particularly in the bound region, where a diverse set of configurations in d_1_ d_2_ space <10-12.5 are quite stable relative to one another. The crystal structure is never found to be the lowest energy state, but we show below that the configurations closest to the crystal structure are among the most stable.

**Figure 4.**
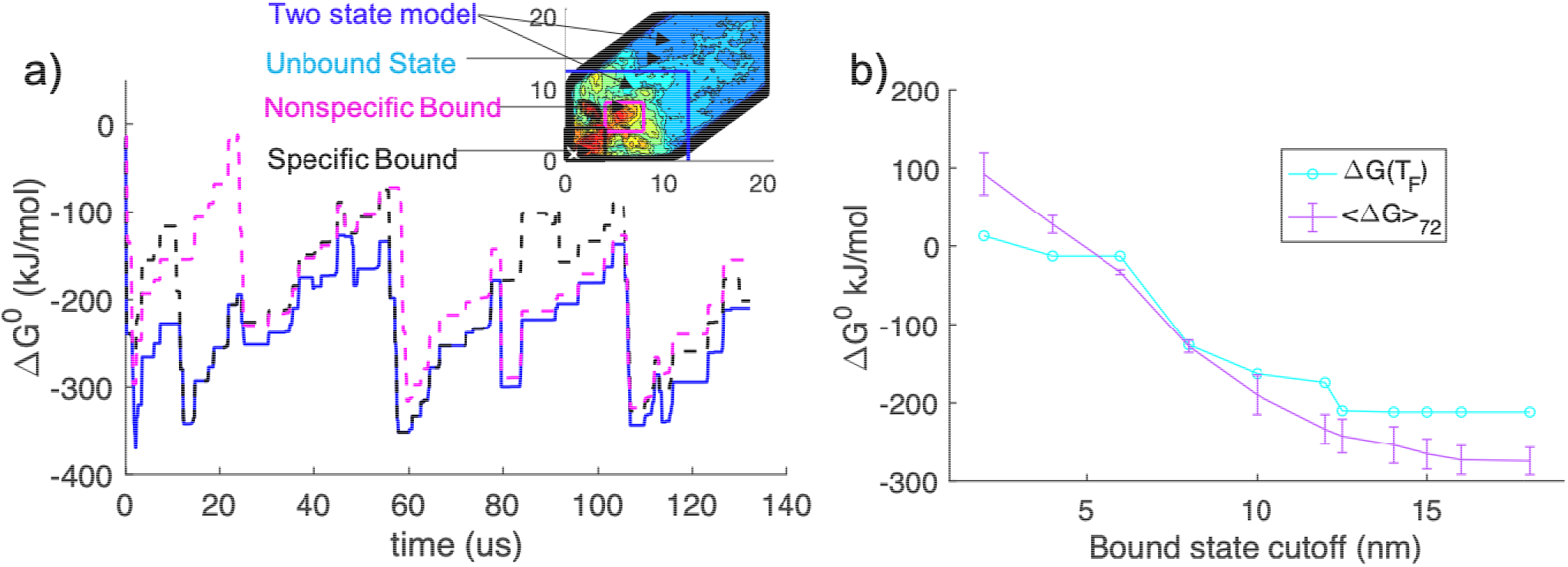
Time evolution of the free energy difference G^0^. a) G^0^ values are reported as bound state free energies relative to the same unbound ensemble, which is always denoted by the region outside the blue lines, where d_1_, d_2_=12.5nm. The time evolution of the blue solid line is the difference between the free energy inside and outside of the blue lines. We define two sub-regions of the bound ensemble for comparison, a specific bound region (black dashed) and a nonspecific region (magenta dashed). The specific bound regions is d_1_<4, d_2_<4.5. The nonspecific region is 4< d_1_<8, and 4< d_2_<8. Significant oscillations in free energies occur due to the relative slow timescale of filling up regions of CV space. The standard deviation of G is 57 (blue), 70 (black) and 54 (magenta) kJ/mol. b) Dependence of the two-state G^0^ on the cutoff in d_1_, d_2_ between bound and unbound. Values averaged after 72μs (purple) and final values (cyan) are shown.

Instead of using the CVs, we can also classify regions of the CV space as bound or unbound based on the properties of the microscopic configurations. Specifically, we can use a distance cutoff between residues to select configurations and their respective grid points, in CV space, for each state. However, this type of metric is problematic, because of the reduced dimensionality of the CV space, where each grid point corresponding to d_1_, d_2_ has an associated degeneracy in terms of the microscopic states. Thus, too many structures with distances greater than a cutoff of ~0.8nm, for example, are found in both the flat unbound region of CV space and in grid points in the stable bound regions. This results in an over-stable estimate of the unbound region, which incorporates grid points in CV space <12nm, where the FE is clearly lower (also see Fig 4b). This indicates that this metric is not effective at discriminating the relative free energies of bound and unbound structures.

### III.C Relative stabilities of specific bound structures

From our free energy surface, the coarse-grained LSP1 proteins clearly form several nonspecific dimers that are quite stable. We perform k-means clustering on our trajectories to define sub-regions of the bound ensemble for structural analysis (Fig 5). We define 7 clusters, with the cluster encompassing the crystal structure region of the surface named as Native. The Central cluster is centrally located on the surface and includes the stable non-specific region. The rest of the clusters are named according to their distinguishing CV value. d_2_4, d_2_8 named according to their d_2_ values while having similar d1 values, d_1_4, d_1_7 named according to their d_1_ values while having similar d_2_ values and d_1_6 named after its d_1_ value. For each cluster, we can then evaluate the thermodynamics of these ensembles. To represent the smaller, more specific crystal structure ensemble, we also evaluated the thermodynamics of configurations with RMSD < 1nm termed as the Crystal cluster. In Fig 6, we show a representative structure for each cluster, chosen by the frame closest to the centroid of each cluster, along with the crystal structure. As described above, we can then quantify the ΔG for each cluster by integrating over that region of the free energy surface, relative to a fixed unbound ensemble defined by the CV space shown in Fig 4 inset.

**Figure 5.**
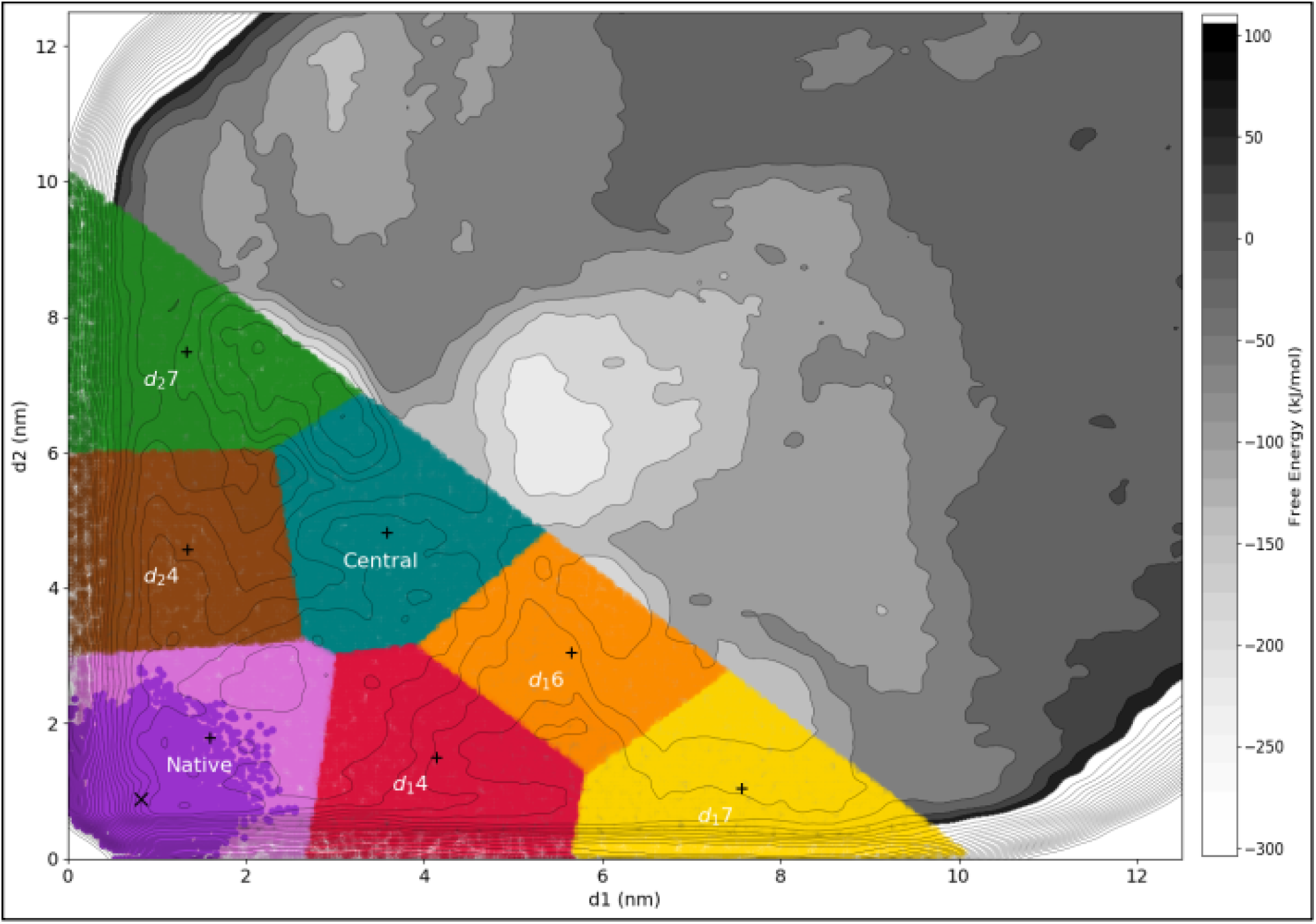
FES after 132 μs with bound ensemble clusters shown. There are seven clusters, configurations of which are projected on an enlarged FES (d_1_<12.5 nm, d_2_<12.5 nm) from Fig 2b. Clustering was performed for all configurations where chain A and B are in contact (see Methods) and with d_1_+d_2_ < 10.2 nm. The crystal structure, denoted by a black X, is included in the native cluster. The darker shade in the native cluster represents the configurations with RMSD < 1nm. Three clusters have relative short values of d_2_, and are then labeled by their d_1_ values via d_1_4 (red; d_1_=4.14, d_2_=1.5), d_1_6 (orange; d_1_=5.66, d_2_=3.0), and d_1_7 (yellow; d_1_=7.57, d_2_=1.04). One cluster has similar d_1_ and d_2_ values, labeled central (teal; d_1_=3.56, d_2_=4.80). Two clusters have short values of d_1_, and are then labeled by their d_2_ values via d_2_4 (brown; d_1_=1.34, d_2_=4.57) and d_2_7 (green; d_1_=1.34, d_2_=7.47). The centroids of each clusters are displayed by a black +.

**Figure 6.**
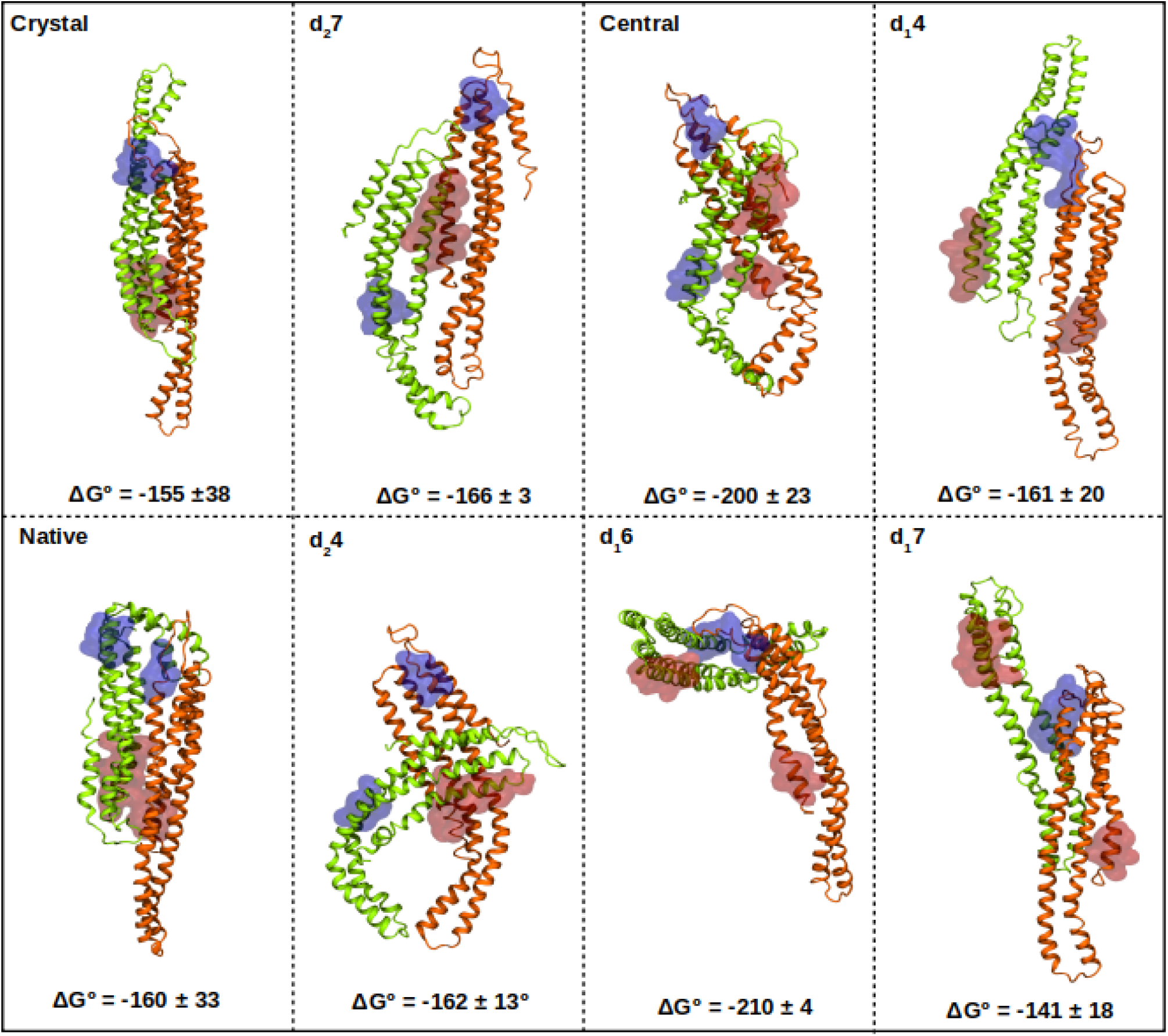
Clustered structures and their associated free energies. Crystal structure contains only a very small region of the CV space around the crystal structure. The other clusters are defined as shown in Fig 5. The native cluster is closest to and encompassing the crystal structure. All Δ*G°* values are in kJ/mol, with SEM. Chain A is in orange and Chain B is green, with the CV patches colored as in Fig 2.

In terms of the potential energy, we do indeed find that the crystal structure has the lowest potential energy of −272 ± 45 kJ/mol. The native cluster has a Δ*U* of −260 ± 22 kJ/mol, which is quite similar to the crystal cluster, consistent with it forming a large and relatively complementary interface. However, two other clusters also have quite stable potential energies, Central and d_1_6, within error of the crystal and native cluster. Central and d_1_6 are adjacent in CV space, with the configurations in Central having a smaller d_1_ value and larger d_2_, and each cluster representing different orientations of the protein chains (Fig 6) as discussed below. These regions separate CV space, with two other cluster sets (d_2_4 and d_2_7) and (d_1_4 and d_1_7), that have clearly weaker potential energies than the native structures (Table 1).

**Table 1.** Thermodynamics of clusters in the bound ensemble

| Cluster | $\Delta G$ (kJ/mol) | $\Delta U$ (kJ/mol) | $-T\Delta S$ (kJ/mol) |
| --- | --- | --- | --- |
| Crystal (RMSD < 1.0 nm) | -155± 38 | -272 ± 45 | 117 |
| Native | -160 ± 33 | -260 ± 22 | 100 |
| Central | -200 ± 23 | -266 ± 56 | 65 |
| d1_6 | -210 ± 4 | -240 ± 10 | 29 |
| d2_7 | -166 ± 3 | -126 ± 46 | -40 |
| d2_4 | -162 ± 13 | -105 ± 66 | -57 |
| d1_4 | -161 ± 20 | -146 ± 21 | -15 |
| d1_7 | -141 ± 18 | -119 ± 10 | -22 |

In terms of the free energy, the native and crystal clusters are not the most stable in pure water, although they are relatively close given the large error estimates, with only d_1_6 appearing to be more stable. The potential energy does not correlate well with the free energy. While this is somewhat unexpected, by characterizing the residue contacts below, we find that there are many residue contacts that form in the nonspecific dimers that can explain the over stability of the bound ensemble for these modeled monomers, and with additional modification to our system including salt, we can see shifts in the relative stabilities. In particular, observing the negative values of entropy differences is unexpected, as they indicate that the bound states have more disorder than the unbound states. However, they are largely within the error, and the unbound ensemble at the standard state concentrations (1M) is also quite limited, with two proteins in a small volume (1.661nm^3^/molecule). The configurations accessible to the unbound ensemble exclude configurations that stabilize nonspecific bound states. We note that the free energies of any cluster in the bound ensemble (Table 1) are not as stable as the full bound state ensemble of −242±22kJ/mol, averaged over the same time period (60-132 *μ*s). This is because at any point in time, the two-state free energy tends to track with whichever basin is lower, and therefore it fluctuates around lower values than any individual basin.

### III.D Native and non-native contacts across the bound ensemble

The native crystal structure BAR domain is stabilized by 50-53 residues per interface, using a 4A cutoff definition. The first two helices from the N terminus (residues 51-136) both have a face that forms the interfacial contacts with the other BAR dimer. The extended tail regions (residues 140-183) do not participate in the binding interface or the membrane binding, but they do clearly form contacts in the filaments observed using cryoEM⍰^4^.The membrane binding residues form patches on the concave surface, R133, K130, R126, R70, K66, K63, R56⍰^2^. The external face of the stable dimer also appears to contribute to stabilization of the filaments formed by the stable dimers⍰^2^.

The LSP1 crystal interface is more hydrophobic than a typical protein-protein interface, which are 30% hydrophobic, 37% charged, and 33% polar^23^. Of the most closely packed residues in the crystal interface, 51% are nonpolar (includes 7% GLY/PRO), 32% are charged, 17% are polar. Thus, only half of the interface is stabilized by polar and charged residues. When the cutoff is increased to include residues out to <6A cutoff, which increases the interface size to 80 residues, we see more hydrophobic residues, such that the total residues are 57% nonpolar. The full protein, which is 214 residues, is 48.6% nonpolar, so there is some enrichment of hydrophobes in the interface, whereas charged drops from 33.2%, polar is at 17.8%. In all regions of the protein ~7% of residues are GLY/PRO. Hence, the overall hydrophobicity of the crystal interface is not significantly higher than other regions of the protein, and as we find, multiple additional surface contacts are quite stable.

We characterized the contacts being formed in each of our nonspecifically bound clusters, as well as our native cluster and the crystal structure, by generating residue level contact maps (Fig 7). The native contacts are highlighted by the diagonal of squares in the top left quadrant of the crystal map (Fig 7 Crystal). Almost all of the contacts are localized in the upper left quadrant, which contains the residues involved in the native interface, partitioned based on whether they are used to define the CVs d_1_, d_2_, or additional regions (d_c_, NP-1, NP-2 see Methods) of the interface. Structures in the native cluster form all of the native contacts present in the crystal structure, with the addition of new contacts between the native patches. There is a shift in the most common contact, which involves native residues off diagonal, although this contact is also present in the crystal structure as well. While there are some contacts formed involving non-native residue contacts (i.e. the lower right quadrant), these are less probable across the cluster.

**Figure 7.**
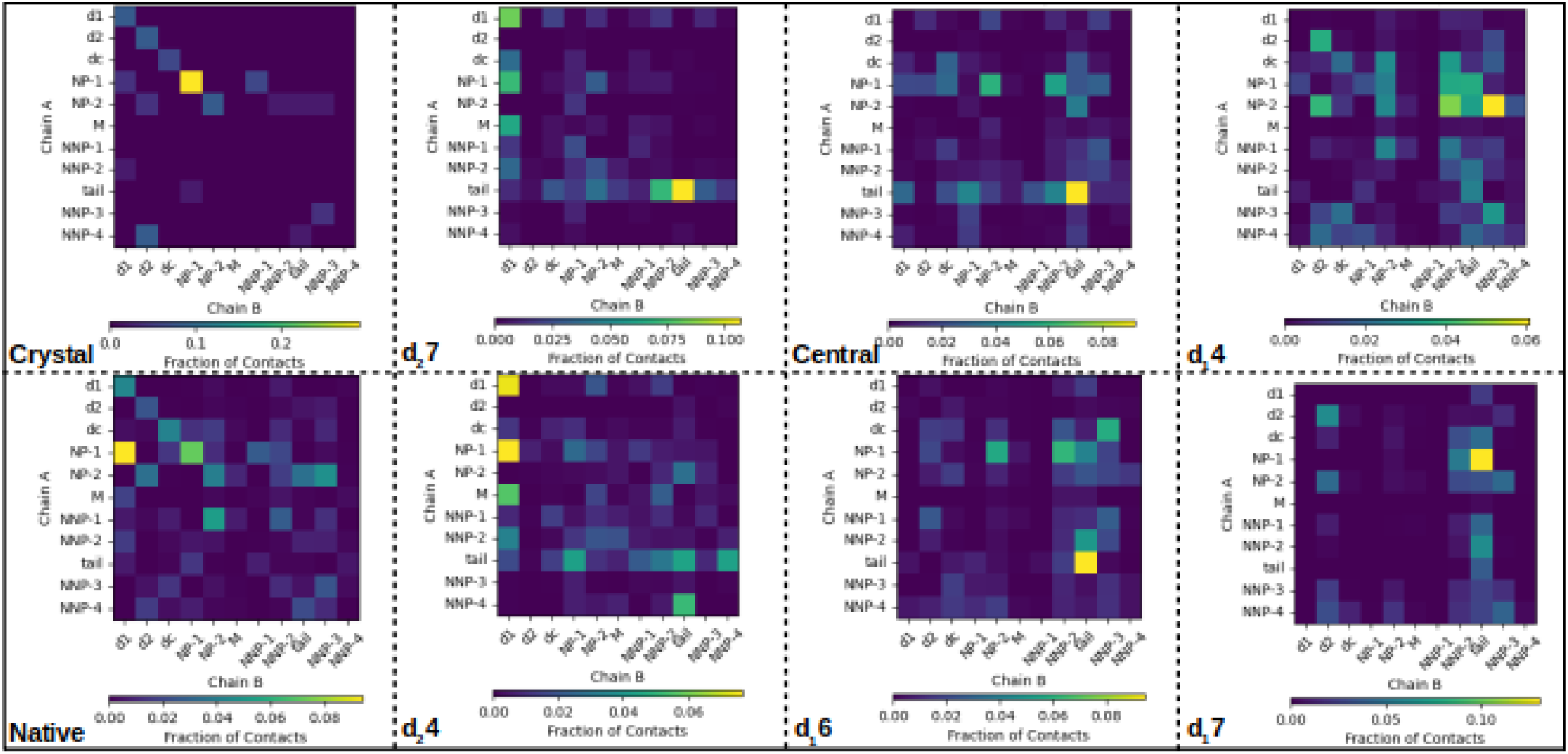
Contact Map. We have partitioned all the residues of the monomer into 11 groups, based on whether they are part of the natively bound interface (d_1_, d_2_, d_c_, NP-1 and NP-2), or residues that are not part of the bound interface (tail, NNP-1, NNP-2, NNP-3, NNP-4). Lastly, the residues that stabilize contacts of the dimer with the membrane are in M. Clusters are the same as in Fig 6. Color bars indicate the fraction of contacts formed between groups, across the ensemble of states per cluster.

A most notable feature of multiple nonspecific dimers is the formation of a strong tail-tail interaction between the chains, which is completely absent in the crystal structure. The tail region juts out from the body of the protein (white region in Fig 1a), defined in the contact map as 35-37 residues (see Methods), with a distribution of 38% nonpolar+PRO, 43% charged, 20% polar. Hence this region is enriched relative to the crystal interface and the full protein in polar and charged residues, which could drive this frequent contact formation. Importantly, the tail regions of LSP1 do form contacts in the higher order filaments based on EM structures^4^, built up from the LSP1 dimers. Hence, the tail regions do participate in specific functional interactions, that are formed simultaneously with the specific crystal interface. Although we do not observe the extended conformation of the LSP1 monomers in this oligomeric conformation, this is perhaps not unexpected when they have yet to form the crystal contacts. Overall, the high probability of this tail region in contacting itself is consistent with these residues forming favorable and ultimately specific contacts in higher-order structures.

As far as the other contacts formed in the nonspecific dimers, d_2_7 and d_2_4, which are fairly close in CV space, form quite similar contacts but with some changes in frequency. The central and d_1_6, which encompass the most stable non-specific dimer forms, are also relatively similar to one another. They lose all of the native contacts in patch d_1_, and instead form highly stable interactions involving the tail regions. The last two clusters, d_1_4 and d_1_7, include contacts between the native d_2_ patches of both chains. Cluster d_1_4 is unusual in that it contains a wide diversity of contacts spread across the surface of the protein, but has a less stable ΔG. This indicates that although it is forming a large number of contacts, they must be significantly weaker interactions.

To simplify the comparison between these clusters further, we quantified the distribution of separations between the dimers (Fig 8a), and the RMSD of each dimer relative to the crystal structure (Fig 8b). The native cluster has a much more narrow distribution of conformations compared to the other clusters, where we see that the central patches of the monomers are always at close separations (Fig 8a). The distribution of RMSD values for the native cluster is closest to the crystal structure, as expected, centered at ~1 nm. In contrast, for the most stable non-specific clusters, central and d_1_6, the RMSD values peak in the range 4-5 nm. The remaining clusters sample a wide range of RMSD values and do not have a single distinctive peak for all the measurements in Fig 8, showing that these clusters include heterogenous sets of conformations.

**Figure 8.**
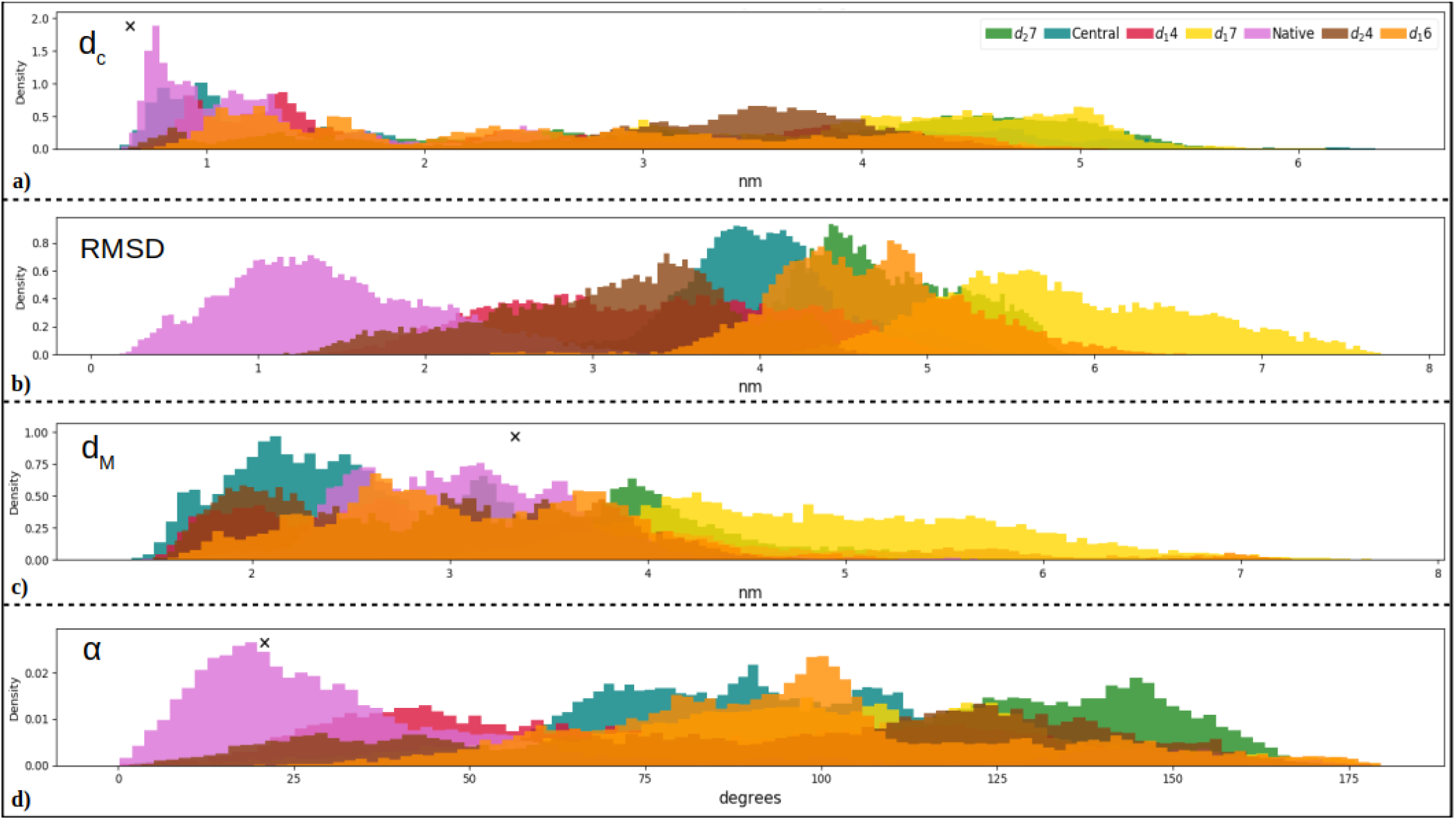
Distance and orientation distributions of bound clusters. The native cluster in mauve encompasses the crystal structure, indicated in black X in all panels, but also has more variation. a) Distance d_c_ tracks the separation between the monomer centers. b) RMSD between chain B and the crystal structure, after aligning chain A (Methods), which is zero by definition for the crystal. c) Distance d_M_ is the separation between the two membrane binding patches (one patch per monomer). d) The angle α reports the orientation of membrane binding patches relative to each other (Methods). In the crystal they are close to a parallel orientation.

### III.E Membrane binding ability in the bound ensemble

The residues involved in binding of the LSP1 dimer to the membrane (Fig 1c) are thus not involved in the specific crystal interface (patch M in Fig 7). Interestingly, these residue patches, which are all positively charged, are almost never involved in the formation of the protein-protein interfaces, even for all of our nonspecific clusters, with only d_2_7 and d_2_4 producing significant contacts between these residues and a set of native residues on the other chain (Fig 7). This indicates that these patches could potentially still bind to the membrane even when in a nonspecific dimer form. We thus evaluated the distribution of distances and orientations between the membrane binding patches in each of our clusters (Fig 8). The distances between the membrane binding patches, d_M_, are broadly sampled in each cluster, and include the separation observed in the crystal structure (Fig 8c). However, the orientation between the two membrane binding patches is quite different between the nonspecific dimers (Fig 8d). Only the native cluster samples orientations that would support simultaneous binding of both LSP1 chains with a membrane surface (Fig 8d). Without proper orientation of the membrane binding patches, the BAR dimer could have inefficient membrane binding and curvature generation. Thus, we expect a strong bias of the native cluster in binding to the surface over any of the other nonspecific dimers, which could form at most one interface with the membrane at a time, rather than two.

### Addition of 100mM NaCl introduces modest changes to the free energy surface

As much of protein-protein association is studied not in pure water but in salty buffer, we ran another set of simulations with the addition of 100mM NaCl. Here we again find a similar stability of the bound structures, including a relatively stable native/crystal structure. Using the same two-state definition as previously, for these shorter simulations we calculate a mean of −224 +− 71 kJ/mol (over two chunks last 60 s), which is comparable to the pure water simulations. While we again find stable nonspecific structures, the overall bound ensemble has contracted, with the stable region now more concentrated along the diagonal of d_1_~d_2_, without extending any further out. We see that the regions defined by clusters d_2_7 and d_1_7 above are now as unfavorable as the unbound state. Most notably, we find that the native structures are the most stable region on the surface (Fig 9a-b). In contrast, in pure water, the nonspecific dimers were of comparable and slightly lower stability than the native structure.

**Figure 9.**
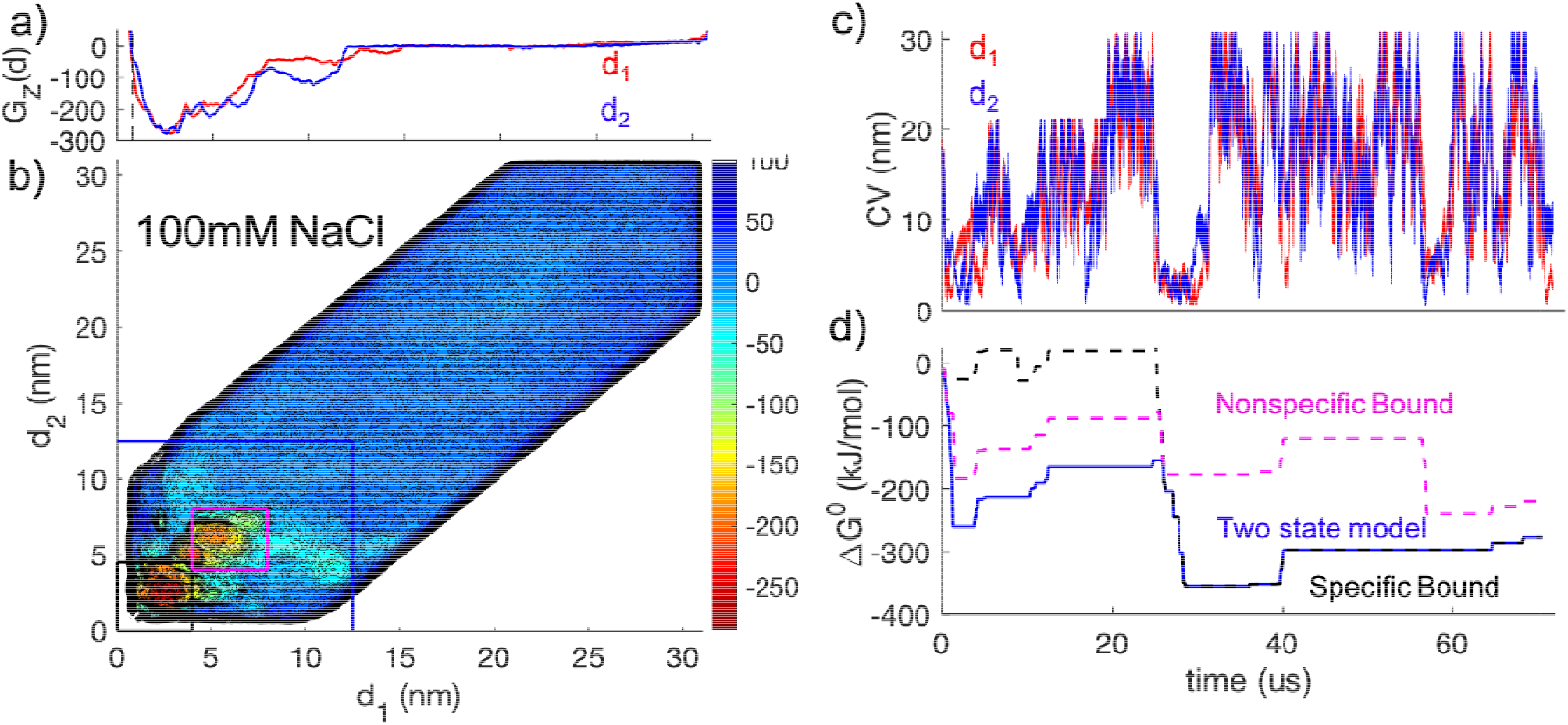
Simulations of LSP1 homodimerization in 100mM NaCl. a) The 1D FES display similar energy scales as without salt, but the nonspecific states are now less, in some cases significantly less stable than the region closest to the crystal (vertical lines). b) The full 2D FES (white X at crystal). All energy units kJ/mol. c) CV sampling in time, where we note we increased the box size from 24nm to 36nm (at 19 s). d) Standard free energy difference as the sampling accumulates shows the specific bound state is determining the overall bound state free energy after 25 s. ΔG at the end of the simulation is −277kJ/mol.

Our results therefore indicate that the addition of salt would lessen the amount of nonspecific aggregation between the LSP1 monomers, and improve formation of the native dimer. We still see nonspecific contacts, especially in the region of clusters d_1_6 and central, which indicate that the tail tail interactions continue to form contacts (Fig 7) with salt present. Although these simulations were shorter, they nonetheless generated multiple transitions in and out of the bound ensemble. They thus provide strong evidence that the solvent can shift the relative stability of these modeled LSP1 dimers, improving selection of the native contacts over non-native contacts.

## IV. Discussion

We find here that the bound state ensemble of coarse-grained LSP1 BAR dimers is highly heterogenous, including a number of nonspecific associations containing large interfaces with typically distinct residue contacts from the native contacts. Nonspecific encounter complexes have been found for a variety of simulated protein-protein association studies^24,31,36^, influencing binding pathways but remaining significantly less stable than the specific bound structure. While these nonspecific structures need to be selected against to prevent aggregation or sequestration of subunits, molecular modeling approaches cannot always accurately predict the proper bound structure, as evidence by continuing efforts to improve prediction methods^61^. Here, we find that the most native-like structure is only the most stable minima when we include 100mM NaCl in solution, otherwise the nonspecific structures are as stable, or slightly more stable. It is significant that the buffer conditions or ionic strength influence the relative stabilities of these structures, as the LSP1 dimer has been crystallized, which would be not be possible if it assembled, as our pure water simulations would suggest, into a heterogenous population of dimers or higher order aggregates. We discuss below how the physiological behavior of the BAR, which requires oligomerization and function on the membrane surface, could drive the heterogeneous bound ensemble. While we find that the native structure is the most stable in salt, the scale of the energy is excessive, comparable to protein folding energies (~90kT) rather than even pM protein association energies that have been the strongest observed affinities for BAR domain dimerization^15^ (~22kT). Two potential origins of this finding: 1) sampling of the bound ensemble 2) force-field limitations and solvent. From our analysis, sampling is not the issue, and instead, modifications to our force-field and solvent choices would improve the energy scale in our modeled LSP1 proteins.

While additional sampling would always be beneficial in MD simulations, our >200μs of metadynamics sampling reproduces, under all simulation conditions and sampling parameters considered, a very strong and heterogenous bound state. The parameters of the metadynamics sampling can impact the final FES, but considering conservative values for both speed of applying bias and variations in height, we do not see any significant differences, indicating our results are not dependent on limited or biased sampling. Because metadynamics ultimately performs a diffusive search in the space of the CVs, there is a time-correlation over which the major basins on the surface will be filled before the surface flattens out again. This effect can be attenuated by dropping the heights of the bias potentials, either manually or using a convergence algorithm^48^, or by applying weighting schemes to reduce the impact of diffusive fluctuations after the surface appears to have sampled the full surface^16^. While lowering our hill heights in particular would help us to reduce large fluctuations in the surface features, as the height exceeds thermal fluctuations, our statistics are sufficient to show that the bound state ensemble is at least 65k_B_T, when averaging of the surface is applied, and more when the average free energy is calculated. The averaged surface predicts slightly smaller free energies of the bound ensemble. This is because each visit to the bound ensemble tends to fill a few basins, not all, and thus on average, they are all less deep. As our movie of the surface construction shows (Mov S1), the fluctuations in the FES in the bound region can also act as temporary barriers for filling up all regions of the bound state, and smaller hills would likely help here. Also, sampling with a third dimension in CV space would provide more paths to reach the native structure and bypass intermediates. Diffusion in 3 dimensions vs 2D will be less sensitive to temporary barriers and any positional dependence of diffusion, which could improve convergence even in a larger space. Overall, we would expect additional sampling may change the relative stabilities of distinct regions of the bound ensemble, as indicated by the error estimates in Table 1, but only modestly, certainly not enough to ‘fix’ the stability of the bound state and push it under 30k_B_T.

The coarse-grained MARTINI force-field is well-studied, accesses much longer-time scales than all-atom MD, and has had significant success in describing biomolecular systems^62^. However, any empirical and coarse-grained force-field does have limitations^63^ that could contribute significantly to the energy scales we observe here for BAR dimerization. Overstabilization and increased barriers between nonbonded interactions^63^ in particular would contribute favorable energy across a large protein-protein interface, relative to the solvated interface. Capturing the native state of a single folded protein chain using MARTINI is already challenging due to the coarsened resolution, and although protein-protein association has lower stability relative to folding, it clearly must be able to select between multiple structurally feasible interfaces. The relative scale of atomic interactions by residue is key, as geometry is not sufficient to select out the native bound structure^36^⍰. Because force-fields are parameterized on properties of small molecule building blocks, modifications can be specifically optimized to the collective behavior of interest, including transmembrane helix dimerization with MARTINI^33^. With a very recent re-parameterization, MARTINI has been shown to effectively capture protein-ligand binding specificity at accurate energy scales^64^. The major re-parameterization changes were in improved packing of the CG beads, and re-parameterized bond distances to improve volume and shape. These changes should similarly improve our characterization of the dimerization of protein-protein interactions, in future work. We also used one of the commonly used elastic network models, ElNedyn, which was not used in the more recent study, and could have contributed to increased flexibility within the monomers that could enable more nonspecific interactions^56^. Consistent with the effect of salt, we found that lowering the dielectric constant from 15 to 1 increased the relative stability of the bound ensemble by a factor 4-5 (data not shown), indicating a sharp dependence on this parameter. Electrostatics, not surprisingly, plays a dramatic role in controlling the relative energy scales of the interface interactions. An increase in the dielectric above 15 is thus one way we could reduce the relative stability of our bound ensemble, with minimal effect on the entropic contributions^28^.

A critical aspect of the force-field is the water model. While the LSP1 specific binding interface is relatively hydrophobic, with ~51% closely packed residues being hydrophobic versus ~30% in an average protein binding interface^23^, the specific interface is not a significantly enriched in hydrophobes relative to the whole protein, which is 48%. The dewetting of each monomer surface to form an interface would benefit other regions on the surface, as well. The ability of water to bridge short range interactions further depends on both its hydrogen bonding ability and size. The size of the MARTINI water is 4 times that of a single molecule, and thus its ability to form favorable interactions at the ~3Å lengthscale are lost, as are a potential gain in stability for interactions bridging the larger water lengthscale in MARTINI. Water-mediated contacts are potentially critical in selection of the native structure relative to nonspecific structures^65^. Polarizability is another critical feature of the water model, where even in all-atom force-fields it has been found to be essential in preventing unphysical aggregation between peptides in water^66^.

Perhaps most interestingly, the physiologic function of the LSP1 BAR domain, and BAR domains in general, is not just to dimerize but assemble into higher-order oligomers, requiring contacts outside of the specific dimer. LSP1 was shown to form ordered filaments even in solution, and were expected to be quite stable assemblies^4^. They do not form nonspecific aggregates, but are more highly ordered, assembling *in vitro* into filaments with a helical tilt^4^, at what we calculate is ~18*μ*M concentration. Our simulations are consistent with this ability of LSP1 monomers to form additional contacts, which would stabilize the dimers assembling into higher-order oligomers, already without the membrane present. LSP1 also has a physiologic partner, PIL1, that is 72% sequence homologous and is even more stable than LSP1 and prone to assembling filaments. PIL1 filaments in solution are not in dynamic exchange with any remaining solution components, indicating that they are nearly irreversible^4^. Of note, PIL1 could not be crystallized, despite having such high homology to LSP1, which could be due to its strong oligomer forming properties. A PIL1 with mutations to membrane binding residues will form filaments in the cytoplasm of cells, rather than in its usual location on the plasma membrane^4^. Further, LSP1 sticks to the plasma membrane containing PIP2, similar to other BAR proteins such as amphiphysin, which will reduce configurational space accessible to the dimers. This would significantly shift the relative stability of nonspecific complexes; as we showed, most of the nonspecific dimers will not be able to align both membrane binding patches with a surface.

Understanding determinants of protein-protein association is essential for building multi-scale models of protein function in cells, where localization and the multi-domain nature of many proteins will impact their function as a population. MD is a powerful tool that, when combined with enhanced sampling, predicts which associated complexes are stable, and which pathways connect the unbound state to the native structure, at resolutions exceeding experiment. Although we cannot extract kinetic rate constants purely from the FES, BD^67^ and MD can be similarly used to extract rate constants^30,37,68,69^, providing a comprehensive picture of binding speeds and intermediates. We find several features of our BAR domain binding pair in common with the still limited number of other protein-protein association studies, including an ability to form nonspecific contacts with barriers between distinct dimer conformations and the importance of electrostatic screening and ionic strength. The domain specific behavior we find here for the BAR domains arises in part because of multiple ‘sticky’ patches on the banana-shaped surface that allow the domain to form both dimers and higher order oligomers. An important application of this approach is that we can apply the same system in the membrane environment, where it functions physiologically. This has the further advantage of minimizing the impact of the force-field parameters, by comparing changes between modeled systems rather than from model to experiment. This will allow us to assess how both the chemistry of the surface, and the restriction to the two-dimensional substrate impact the bound ensemble and the dissociation constant. While membrane curvature changes are a critical feature of BAR domain function, the dissociation constant on the membrane surface can only be roughly estimated from the 3D value^70^. Details on entropic and enthalpic changes to the bound ensemble on the surface are needed to determine the 2D dissociation constant^70,71^. This value is an important ingredient in models of BAR domain behavior as a population in the cell, where localization to the 2D surface can dramatically increase the formation of assemblies via dimensional reduction^72^. Our results provide an important benchmark for the study of dimerization in large protein-protein association, adding in the first representative from the critical class of membrane sculpting proteins.

## ACKNOWLEDGEMENTS

We acknowledge support from the MARCC supercomputing center at JHU. MEJ gratefully acknowledges support from an NIH MIRA award R35GM133644.

